# Zinc Solubilizing Plant Growth Promoting Microbes Produce Zinc Nanoparticles

**DOI:** 10.1101/602219

**Authors:** Uzma Sultana, Suseelendra Desai, Gopal Reddy, TNVKV Prasad

## Abstract

Strains of *Pseudomonas, Bacillus* and *Azospirillum* with plant growth promoting ability were checked for their zinc solubilizing ability. Efficient zinc solubilizers were checked for their ability to produce nano-scale zinc particles. The nanoparticles from the cell-free culture filtrates obtained from these strains were characterized for particle size, Zeta potential and functional groups. Presence of Zn nanoparticles in the bacterial culture filtrate was confirmed by particle distribution and Scanning electron microscope (SEM) analysis. Most properties of nanoparticles are size dependent. Zinc nanoparticles were observed to be spherical in shape and size ranged from 52.0 to 106.0 nm. Zeta potential of the Zn nanoparticles was estimated to understand the stability of the particles. The measured zeta potentials varied from −14.5mV to +179.10 mV indicating high stability and dispersion of the zinc nanoparticles. FTIR peaks at different wave numbers depicted the role of functional groups of proteins in the biosynthesis of Zn nanoparticles. This finding opens a new area of research focusing on microbe-microbe interactions in rhizosphere and plant-microbe interactions at rhizosphere apart from biosynthesis of nanoparticles, which has major applications. To our knowledge, this is the first report of production of nanoparticles as part of nutrient mobilization by plant growth promoting rhizobacteria.

## 1. Introduction

Food- and nutritional-security are major concerns of most of the developing nations. Production of sufficient and nutritious food is a major challenge in front of many countries including India. Zinc (Zn) is one of the vital minerals required for optimum plant growth as well as animals, including human beings such as mental retardations, impairments of the immune system and overall poor health. In recent years, the Zn deficiency problem has received increasing attention and appears to be the most serious micronutrient deficiency together with vitamin A deficiency. Zinc deficiency is particularly widespread among children and represents a major cause of child death world over. It plays a vital role in biochemical reactions in plants and its deficiency is displayed as a remarkable reduction in plant height and develops whitish-brown patches that turn necrotic subsequently (Hughes and Poole;1989). Zn occurs in soil as sphalerite, olivine, hornblende, augite and biotite and availability of Zn from these sources is guided by many factors among which bio-chemical actions of rhizo microorganisms play an important role (Bhupinder Singh 2005). The solubility of zinc is highly dependent upon soil pH and moisture and this is the reason why arid and semi-arid areas are zinc deficient. Most of the Zn in soil is in unavailable form and thus unable to meet nutritional requirements of plants. In India, >50% of agro-ecosystems are Zn deficient, especially the Indo-Gangetic plains spanning across the States of Punjab and Uttar Pradesh. Overall, zinc deficiency in the country has increased from 42% to 49% in the past four decades and it is expected to increase up to 63% by 2025 (Singh 2009). Plant response to Zn deficiency occurs in terms of membrane integrity loss, reduced synthesis of metabolites and susceptibility to heat stress (Bhupinder Singh 2005). Zn is supplemented exogenously to plants in the form of chemical fertilizers like zinc sulphate that subsequently transforms (96-99%) into different insoluble forms depending upon the soil types, physico-chemical reactions and thus becomes totally unavailable in the environment within seven days of application (Rattan and Shukla 1991; Venkatakrishnan 2003).

External supplementation of micronutrients is not only expensive, but also leads to unwanted accumulation of them in the soil in fixed forms. Accumulation of excess Zn complex leads to deterioration of soil-health, integrity and microbial diversity. Further, at a given point of time, plants need only small quantities of micronutrients. Efforts to supplement nutrients in the form of biofertilizers helped the rainfed farmers significantly (Venkateswarlu and Wani 1999). Strains of species of *Pseudomonas, Bacillus, Azotobacter, Azospirillum* and *Acetobacter* etc. are known to promote plant growth, (Kloepper 1991 and Orhan 2006). Solubilization of mineral nutrients; stimulation of root growth; and suppression of root diseases are some of the modes of actions of plant growth promoting rhizobacteria in influencing plant health and productivity (Martinez-Viveros 2010). Bacteria-mediated solubilization of phosphorus (P) to supplement P requirement of the plants is a popular technology among farmers in many countries (Cattelan 1999; Gull et al., 2004; Richardson et al 2009; Leo Daniel Amalraj et al., 2014). Similarly, Zn solubilization by microorganisms has been widely studied in fungi and bacteria (White et al., 1997; Di Simine et al., 1998; Fasim et al., 2002; Suseelendra Desai et al., 2012). Nanoparticles with small size and large surface area are expected to be the ideal candidates for use as a Zn fertilizer in plants (Prasad 2012). Particle size may affect the agronomic effectiveness of Zn fertilizers. Decreased particle size results in increased number of particles per unit weight of applied Zn. Decreased particle size also increases the specific surface area of a fertilizer, which should increase the dissolution rate of fertilizers with low solubility in water, such as zinc oxide (ZnO) (Mortvedt, 1992). Nanoparticles of Mg, Zn, Fe and P are the structural component of enzymes (phosphatases and phytase), polysaccharides and chlorophyll. In plants, nanoparticles can be applied for a broad range of uses, particularly to tackle Phytopathological infections, nutrition supplement and as growth adjuvant. These nanoparticles can be tagged to agrochemicals or other substances as delivery agent to plant system and tissues for controlled release of chemicals. They are known to stabilize the enzyme complexes in plants. First evidence of biosynthesis of nanoparticles was reported using *Pseudomonas stulzeri* where the nanoparticles were deposited on the cell membrane (Klaus 1999). Subsequently, *Bacillus licheniformis* (Kalimuthu 2008); *Lactobacillus* strains (Nair 2002); *Bacillus subtilis* (Saifuddin 2009); *Cornybacterium* sp. (Zhang 2005); and *E.coli* (Gurunathan 2009a; Gurunathan 2009b) were shown to be involved in extra and intracellular synthesis of nanoparticles. (Minaeian 2008) reported synthesis of silver 50-100 nm particles by *Klebsiella pneumoniae*, and *Escherichia coli*. While a number of reports are available on the biological synthesis of silver nanoparticles by potential endophytic microorganisms, scanty information is reported on the synthesis of Zn nanoparticles using plant growth promoting bacteria. In the present study, we report the ability of Zn solubilising rhizobacteria isolated from various crop production systems of different agro-ecological sub regions of India to produce Zn nanoparticles.

## 2. Materials and methods

Bacterial strains used in this study were obtained from the culture bank of ICAR-Central Research Institute for Dryland Agriculture, Hyderabad. Based on previous qualitative and quantitative screening, five efficient zinc solubilising strains viz. *Bacillus* B116, *Pseudomonas* P29, *Pseudomonas* P33, and *Azospirillum* 20 were selected for nanoparticle analysis (Praveen Kumar 2012).

All the isolates were grown in Minimal salt medium amended with 0.1% ZnO with an initial population containing 2×10^8^ CFU/ml and incubated at room temperature in an orbital shaking incubator with 180 rpm speed. The culture medium was centrifuged after 72h of inoculation and the supernatant was collected. Supernatant was analysed for the presence of nanoparticles in nanoparticle analyzer.

### 2.1 Characterization of Zn nanoparticles

Ten mL of the supernatant was transferred to a cuvette and placed into the nanoparticle analyzer (Horiba Nanoparticula 100). Three replicas for each sample were maintained respectively. Each sample was read three times and the mean was considered as particle size. The zeta potentials were recorded using Particle size analyzer (Horiba Nanoparticula 100, Japan). This apparatus uses Dynamic Light scattering (DLS) phenomenon. The sample holder temperature was maintained at 25°C, a technique used to determine the stability profile of small particles in solution. Nano scale Zinc particles were observed under SEM. To observe nanoparticles of Zinc, 10 μl of the suspension was spotted on to the metal slide. Metal slide was placed inside a desiccator over night to remove moisture completely. After removing the slide from the desiccator, a gold sputtering of approximately 1nm was done over the suspension.

For Fourier Transform Infra Red (FTIR) spectroscopy measurement, the bio-transformed product present in cell-free filtrate was analyzed. FTIR spectrum (Bruker Tensor 27, USA) was recorded in mid IR region in the range of 400–4000 wavenumber cm^-1^. A drop of the sample solution was carefully added to the potassium bromide crystals using a micropipette and the spectrum was recorded in transmittance (%) mode.

## 3. Results

The results of the particle size distribution (PSD) dynamic light scattering method conforms the presence of Zinc nanoparticles. Particle Z-Average size and Poly dispersity index are presented in table 1.

**Table 1.** Particle Z-Average size and Poly dispersity index of Zn nanoparticles produced by different bacterial strains in ZnO incorporated broth.

| Strains | Z-Average particle size<br>(nm) | Poly dispersity Index |
| --- | --- | --- |
| B116 | 593 .0 | 0.60 |
| P 29 | 428.8 | 0.51 |
| P 33 | 482.0 | 0.37 |
| Asp 20 | 539.0 | 0.48 |
| Control | 6279 | 1.00 |

Electron Microscopy (SEM) analysis image provided further insight into the shape size, morphology and distribution of nanoparticles. The SEM micrographs recorded, showed spherical or roughly spherical nanoparticles, which were observed to be uniformly distributed. The drop coated film of the Zn nanoparticles were distributed on the surface and were uniformly dispersed with some aggregation. The size of the Zn nanoparticles ranged from 106 to 52nm. In control the size was 5200nm (Table 2 and Fig 1).

**Table 2.** Size of available Zn (nm) in MS medium amended with ZnO.

|  |  |
| --- | --- |
| Minimal salt media | Size (nm) |
| amended with ZnO |  |
| Control | 5200 |
| B 116 | 106 |
| P 29 | 52 |
| P 33 | 74 |
| Asp 20 | 53.4 |

**Fig 1.**
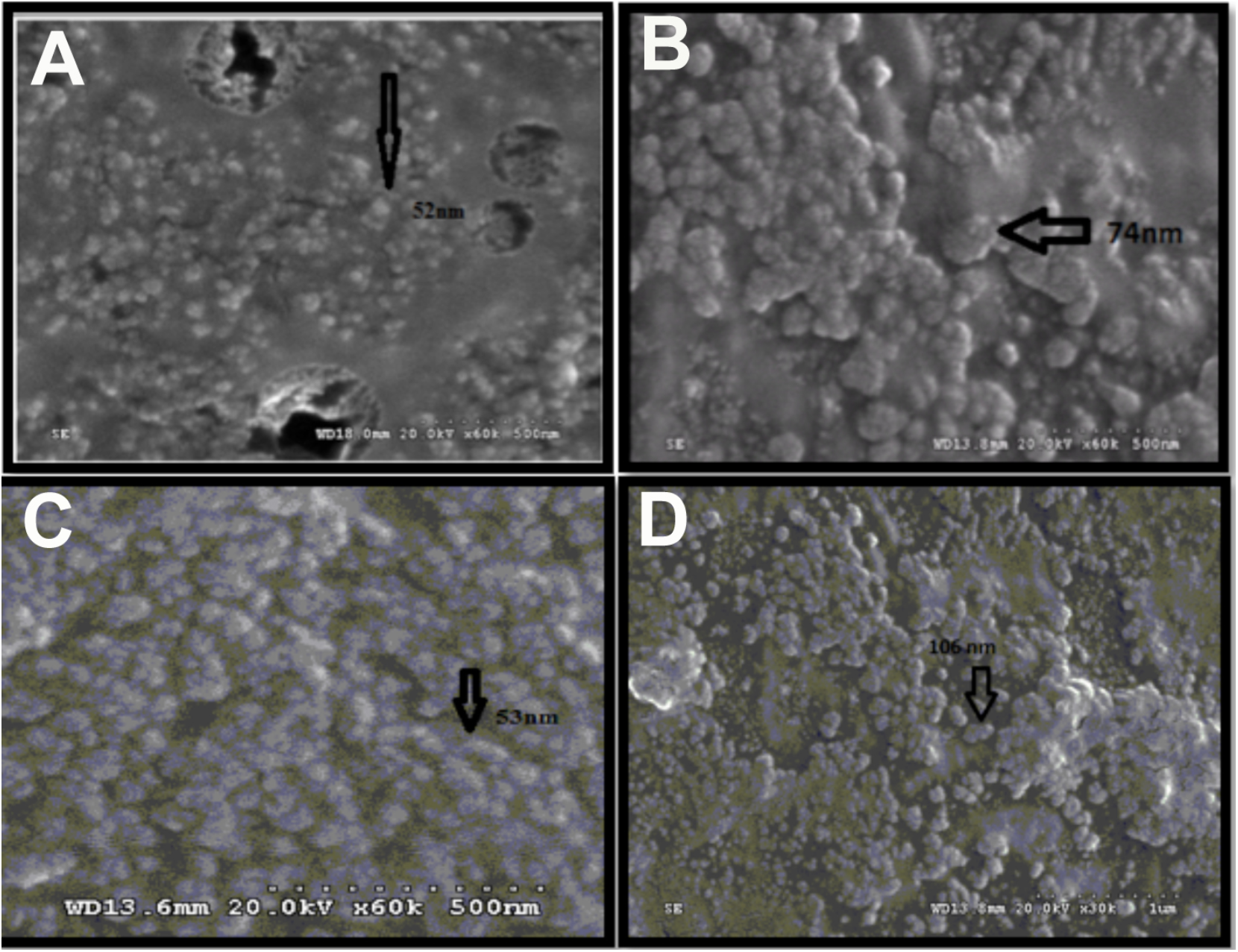
SEM image of Nanoparticles of zn produced by a) P 29 (52nm) b) P 33 (74nm) c) Asp 20 (53nm) d) B 116 (106nm) in media amended with 1ppm ZnO.

### 3.1 Zeta Potential

Using the nano particle analyzer, the Zeta potential was measured as the magnitude of ZetaPotential (−200mV to +200mV) gives an indication of the potential stability of the colloidal system. The Zeta potentials of Zn nanoparticles produced by B116, P33, P29 and Asp20 were +179.1 mV, −14.5mV, + 0.5mV and 0.5 mV respectively, clearly indicating the stability of the nanoparticles (Fig 2).

**Fig 2.**
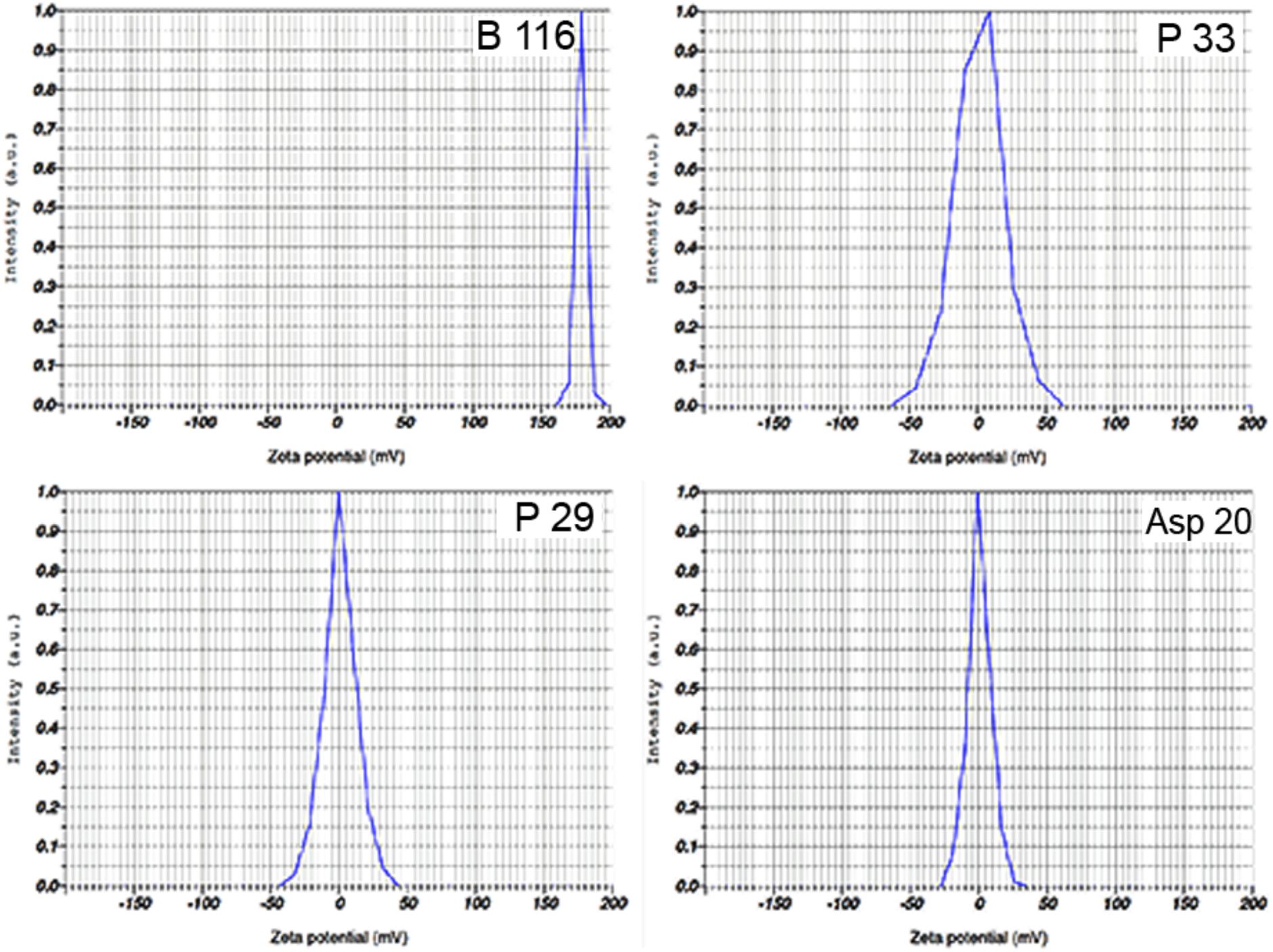
Zeta potential of Zn nanoparticles produced by B116 (+179.1 mV), P33 (−14.5 mV), P29 (+0.5 mV), Asp 20 (+0.8 mV), and in ZnO amended MS medium

### 3.2 FTIR Analysis

FTIR analysis of the cell-free culture filtrates of ZnO amended MS medium was done to identify the interactions between Zn and bioactive molecules enabling synthesis and stabilization of Zn nanoparticles (Fig 3). The FTIR spectra of cell-free extracts of ZnO amended MS medium showed prominent peaks at 3346, 2923, 1636, 1459 and 538 wave numbers cm^-1^. The strong broad wave at 3346 cm^-1^ is attributed to NH stretch of amides. The wave observed at 2923 cm^-1^ is stretch vibration of primary and secondary amides of protein. The band observed at 1636 cm^-1^ is assigned to the bending vibrational mode of amides. The peaks at 1459 refer to amino and amino-methyl stretching groups of protein. Peak observed at 529cm-^1^ corresponds to the stretching vibrations of Zn nanoparticles.

**Fig 3.**
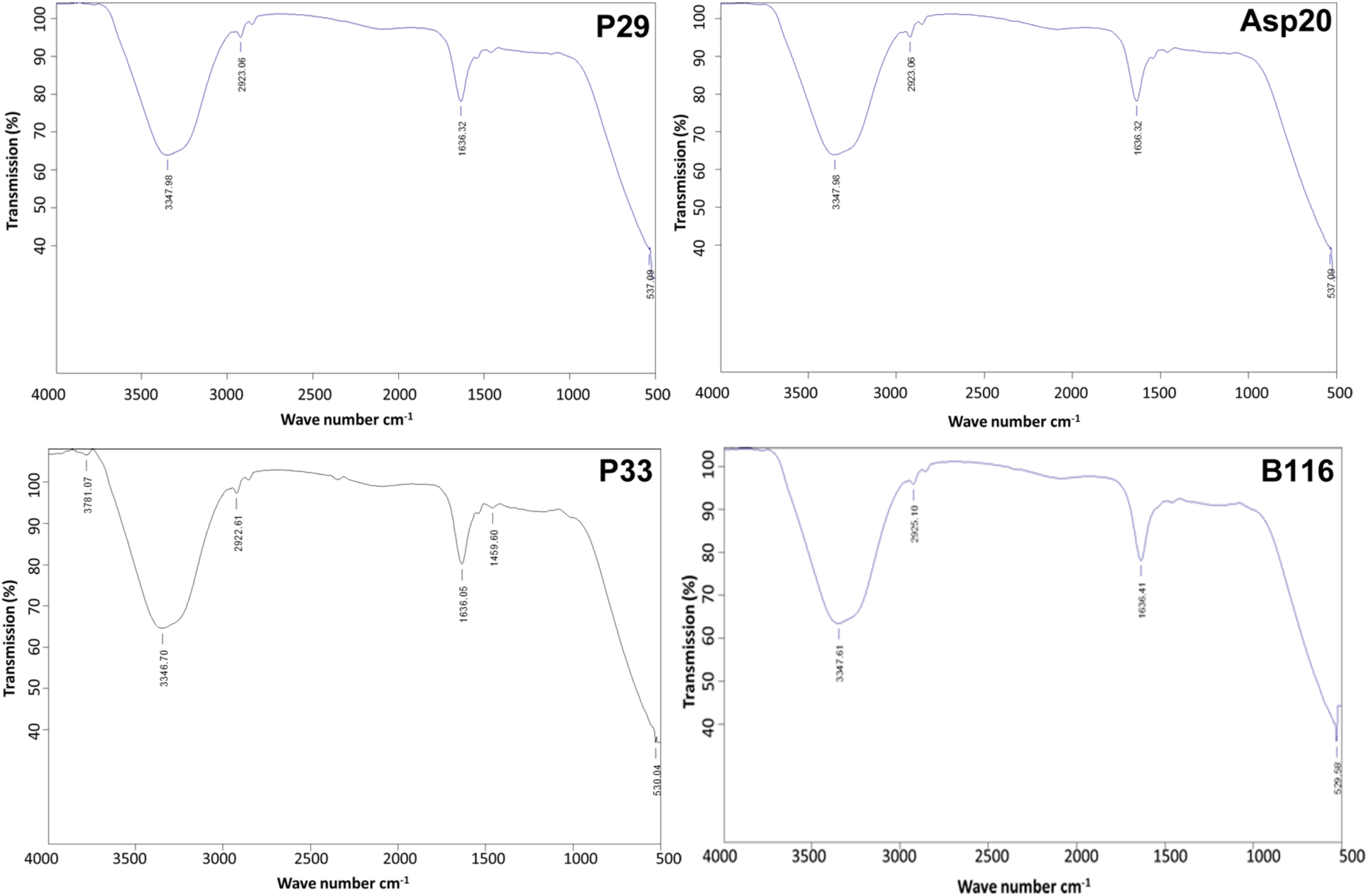
IR spectrum of cell-free culture filtrates of ZnO amended MS medium inoculated with different strains of bacteria

## 4. Discussion

Plant growth promoting strains of *Pseudomonas, Bacillus* and *Azospirillum* found to solubilize ZnO and ZnCO_3_ were formulated and field-evaluated. While trying to understand their mode of action of Zn solubilization, we found that these strains synthesized Zn nanoparticles *in vitro* when cultured in MS broth supplemented with ZnO or ZnCO_3_. There were differences among the strains in synthesizing different sizes of zinc nanoparticles. According to particle size distribution and SEM analysis, *Pseudomonas putida* P29 produced smallest zinc nanoparticles of the size 52 nm followed by Asp20 and B116 produced large particles (106 nm) as against control where the particle size was 5200 nm. The exact mechanism for the synthesis of nanoparticles using biological agents has not yet been elucidated but it has been suggested that various biomolecules are responsible for the synthesis of nanoparticles. The mechanism of extracellular biosynthesis of nanoparticles is proposed as a reductase-mediated reaction, which brings about bioreduction of metal ions and synthesizes nanoparticles (Honary et al., 2012). In the present study, FTIR spectrum of the nanoparticles indicated the presence of various chemical groups, one of which is an amide. An amide I band was observed at 1636 wavecm^-1^, which is the bending vibrational mode of amides. The stretching band of amide at 3346 wavecm^-1^ further confirmed this result. The 1636 wave cm^-1^ corresponds to amide I due to carbonyl stretch in proteins. FTIR spectrum showed the presence of functional groups such as amide linkages and –COO–, possibly between amino acid residues in protein and the synthesized Zn nanoparticles. This report is in line with the findings of (Honary 2012) which support the interactions between the proteins and nanoparticles during the biosynthesis. The surface properties of the nanoparticles are known to be one of the most important factors that govern their stability and mobility as colloidal suspensions, or their adsorption or aggregation and deposition. Zeta potential can be related to the stability of colloidal dispersions (Vielkind 2013) as it indicates the degree of repulsion between adjacent and similarly charged particles in dispersion and high zeta potential values indicate high stability and with decrease in the particle size, the stability also decreases. Highest zeta potential of +179.10 mV was observed for the particle size 106.0 nm produced by B116 indicating best stability. However, nanoparticles with zeta potential values greater than +25mV and −25mV typically are stable (Sonika 2015). Hence, it could be inferred that the nanoparticles produced with lower zeta potentials such as +0.5mV may not be stable even though they are only 52 nm. Further, low stability nanoparticles tend to form aggregates in the medium, which is influenced by various factors. Aggregation of metal oxide nanoparticles in aqueous solutions depends on ionic strenght, pH, surface charge and charge density (Guzman 2006; Domingos 2009). In the present study also, such aggregation was noticed in cases where the zeta potential was low. Results of particle size distribution clearly indicated that the obtained Zinc nanoparticles are mono as well as poly dispersed in nature. Samples with very broad size distribution have Poly dispersity index values >0.7. These findings are supported by the findings of (Anil reddy 2009).

## 5. Conclusion

Plant nutrient management is an important domain of research and needs understanding of the basic phenomena of uptake and utilization of all essential nutrients. Zn is an essential nutrient which has a major role in several plant- and human-metabolic pathways. The present finding opens up a new area of research of not only green synthesis of nanoparticles using biological systems but also the dynamics of inter-organism nutrient dependence in the rhizosphere. In this study, we have reported for the first time synthesis of Zn nanoparticles by the plant growth promoting bacteria with a particle size ranging from 52 to 106 nm. Further research is underway to understand role of these Zn nanoparticles as a factor in rhizosphere dynamics and modulation, its uptake and transport in crop systems for increasing the productivity.

## Acknowledgements

The authors sincerely thank Director, ICAR-CRIDA for his constant support and encouragement to conduct this research at CRIDA. Senior author also acknowledges the fellowship from University Grants Commission (MANF scheme) and Dr. B. Suresh ARCI Hyderabad for providing research facilities.

